# Functional and evolutionary insights into the emerging *tet*(X4)-carrying non-O1/O139 *Vibrio cholerae* from retail pork

**DOI:** 10.64898/2026.08.12.744420

**Authors:** Mengjia Hui, Xinyi Huang, Bingbing Li, Furong Ding, Xiang Liao, Haoqing Lu, Xing Shi, Liurui Liang, Kaichao Chen, Xiaofan Li, Huimin Si, Chen Xu, Ping Zeng, Sheng Chen, Ning Dong, Qipeng Cheng

## Abstract

The tigecycline resistance gene *tet*(X4) is prevalent in Enterobacteriaceae, particularly in *Escherichia coli*. To our knowledge, no study has reported the dissemination dynamics of *tet*(X4) in *Vibrio* spp. Herein, we isolated and characterized a first *tet*(X4)-positive non-O1/O139 *Vibrio cholerae* isolate from retail pork. Genomic sequencing identified a novel *tet*(X4) variant in the *V. cholerae* chromosome, harboring a G568A nucleotide substitution that resulted in an Ala190Thr (A190T) amino acid substitution in Tet(X4). While this Tet(X4)-A190T variant conferred lower phenotypic resistance to tetracyclines (including tigecycline) than the wild-type Tet(X4), its overall catalytic efficiency against these antibiotics was paradoxically enhanced despite a reduced substrate affinity. Genomic comparisons revealed that two copies of IS*CR2* flanked the variant gene, and the structure was IS*CR2*-*hp*-*hp*-*abh*-*tet*(X4)^G568A^ -IS*CR2*, which is highly homologous to the reported *E. coli* plasmids carrying *tet*(X4). In addition, it confirmed the presence of an IS*CR2*-mediated circular intermediate, proving this module’s capacity for horizontal transfer of the *tet*(X4)^G568A^ variant. Furthermore, the IS*CR2*–*tet*(X4) genetic structure carrying the G568A substitution was integrated within a chimeric SXT/R391-like integrative and conjugative element (ICE), which is also serving as a vehicle for genetic dissemination. As per our knowledge, this is the first report on the emergence of SXT/R391-like ICE carrying *tet*(X4) in *Vibrio* strains. Our finding demonstrates that the clinically relevant tigecycline resistance gene *tet*(X4), previously confined mainly to Enterobacterales from humans and livestock, is now actively spreading into environmental *Vibrio* populations. This cross-species transfer highlights a previously underappreciated ecological and public health concern in aquatic ecosystems.

**Importance:** Tigecycline serves as a vital last-resort antibiotic against severe multidrug-resistant bacterial infections, but its clinical efficacy is currently threatened by the rapid global dissemination of resistance genes like *tet*(X4). While land-based agriculture is a well-recognized reservoir for these genes, the role of aquatic ecosystems and environmental pathogens, such as *V. cholerae*, in harboring *tet*(X) determinants remains largely unexplored. In this study, we characterize a non-O1/non-O139 *V. cholerae* isolate from retail pork that harbors a naturally occurring, chromosomally integrated *tet*(X4)^G568A^ variant. This novel variant exhibits elevated catalytic efficiency against tetracycline antibiotics. The *tet*(X4)^G568A^ allele is embedded in a highly conserved structural module (IS*CR2*–*tet*(X4)–*abh*–*hp*–*hp-*IS*CR2*) flanked by two IS*CR2* repeats, which is integrated into an SXT/R391-like ICE at the chromosomal *prfC* locus. These findings provide the first high-confidence genomic evidence of *tet*(X4) in *V. cholerae*, highlighting aquatic *Vibrio* species as critical environmental reservoirs for clinically significant antimicrobial resistance genes and emphasizing the urgent need for continuous genomic surveillance.

## Observation

Tigecycline has served as a critical last-resort antibiotic for infections caused by multidrug-resistant Gram-negative pathogens(1, 2). However, the recent emergence of plasmid-encoded resistance genes such as the *tet*(X) variants(3, 4) poses a severe threat to the continued clinical efficacy of tigecycline. Among them, the *tet*(X4) variant is exceptionally significant, driven by highly efficient mobile genetic elements such as IS*1*, IS*26*, and particularly IS*CR2*(3, 5, 6), it has successfully breached species barriers and established broad ecological dissemination, particularly in farming and food production chains(7-9).

While land-based livestock farming is a recognized reservoir for antimicrobial resistance genes (ARGs), aquatic ecosystems play an equally critical role in their global spread. *Vibrio* spp. are typical environmental bacteria distributed in estuaries and coastal waters(10), are increasingly recognized as vital environmental reservoirs and cross-species transmission vectors for harboring diverse ARGs(11, 12). However, the presence and functional role of *tet*(X) genes in *Vibrio* species have remained largely unexplored.

Here, we identified the first *tet*(X4)-positive non-O1/non-O139 *V. cholerae* isolate (WH-V113) from retail pork during surveillance of foodborne *Vibrio* **(Table S1)**. Whole-genome sequencing revealed a naturally occurring G568A mutation in the *tet*(X4) gene of this isolate, which encodes a Tet(X4) protein with a single amino acid substitution (A190T) at residue 190, replacing alanine with threonine **(Fig. 1A)**. Antimicrobial susceptibility testing revealed that WH-V113 exhibits a multidrug-resistant phenotype. Specifically, the strain was resistant to tetracycline (MIC = 16 mg/L) and tigecycline (MIC = 1 mg/L), while maintaining low MICs for doxycycline (2 mg/L), omadacycline (2 mg/L), eravacycline (1 mg/L), and minocycline (<0.12 mg/L) **(Fig. 1B)**. Additionally, WH-V113 demonstrated resistance to ampicillin, cefotaxime, trimethoprim-sulfamethoxazole, and colistin, but remained susceptible to amikacin, ciprofloxacin, gentamicin, meropenem, and azithromycin **(Table S2)**

**Figure 1.**
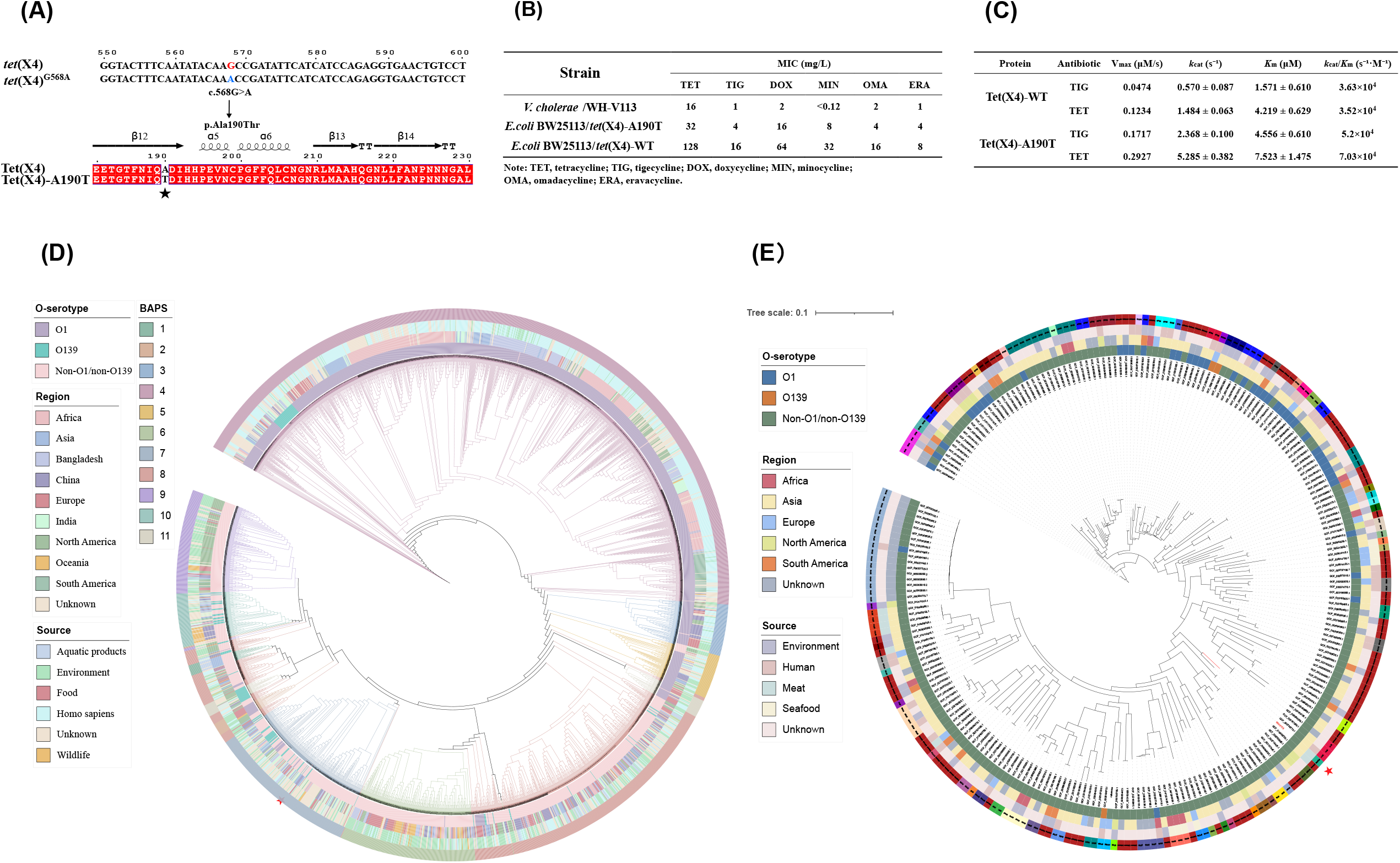
Function and Phylogenetic analysis of *tet*(X4)^G540A^ variant non-O1/non-O139 *V. cholerae* isolate. **(A)** Sequence alignment *tet*(X4)^G540A^ variant with wild type;**(B)** MIC values of tetracycline antibiotics against *tet*(X4) carrying strains. **(C)** Kinetic parameters of wild type Tet(X4) and Tet(X4)-A190T variant toward tetracycline and tigecycline. **(D)** A maximum-likelihood phylogenetic tree was constructed based on single-nucleotide polymorphisms (SNPs) using Snippy and IQ-TREE. The analysis included all *V. cholerae* genomes downloaded from the NCBI database (accessed on April 23, 2026) and the WH-V113 isolate. The genomes were grouped by country of origin. The outer rings represent: **(1)** O-serotype, **(2)** geographical origin (country/region), **(3)** sample source, **(4)** BAPS cluster. Detailed information corresponding to the color codes in rings 1–3 is provided in the legend. **(E)** A focused SNP-based phylogenetic tree was reconstructed using all isolates assigned to the same fastBAPS cluster as WH-V113 (highlighted in Figure 2A). The outer rings represent: **(1)** O-serotype, **(2)** geographical origin (country), **(3)** sample source, and **(4)** MLST profile. The specific MLST profiles are explicitly labeled on the outer ring, while detailed color-coding information for the other rings can be found in the legend above.

To explore the functional effects of A190T substitution, antimicrobial susceptibility testing was performed using *E. coli* BW25113 expressing either the wild-type Tet(X4) or the Tet(X4)-A190T variant. The Tet(X4)-A190T variant-expressing strains exhibited lower MICs for most tetracyclines compared to those expressing the wild-type **(Fig. 1B)**, indicating an attenuated resistance phenotype. However, subsequent enzyme kinetic analyses revealed a surprising paradox: Tet(X4)-A190T variant possessed higher *k*_*cat*_, *K*_m_, catalytic efficiency (*k*_*ca*t_/*K*_*m*_), and Vmax values toward both tetracycline and tigecycline than the wild-type enzyme **(Fig. 1C, S1)**. Specifically, the catalytic efficiency increased approximately 2.0-fold for tetracycline and 1.4-fold for tigecycline. Structural predictions indicated that the overall protein architecture remained highly conserved with wild-type(13) (RMSD = 0.347 Å) **(Fig. S2)**. The introduction of a polar hydroxyl group in β-sheet 12 (β12) likely perturbs the local microenvironment surrounding the FAD binding pocket, modulating the redox potential of the flavin cofactor and increasing overall catalytic turnover despite reduced substrate affinity. The integration of these findings reveals a significant discrepancy between enzymatic activity and phenotypic resistance. While nearly all reported *tet*(X4)-positive isolates exhibit a tigecycline MIC of ≥8 mg/L(3, 4, 14-16), the WH-V113 isolate demonstrated an unusually low MIC of 1 mg/L. Yet, heterologous expression of this variant in *E. coli* yielded 2∼16 folds increase in tetracycline MICs compared to the parental WH-V113 strain, paired with markedly increased catalytic efficiency *in vitro* **(Fig. 1B)**.This discrepancy strongly suggests that catalytic efficiency alone does not dictate *tet*(X4)-mediated resistance in *V. cholerae* WH-V113 and that additional, uncharacterized host-related factors are likely involved in conferring high-level resistance, necessitating further exploration.

Genomic characterization of WH-V113 further expanded our understanding of *tet*(X4) acquisition and stabilization. The complete genome consisted of a primary chromosome (Chromosome I, 3088970 bp) and a secondary chromosome (Chromosome II, 1030151 bp), with no plasmids detected (Fig. S3). Multilocus sequence typing (MLST) revealed that strain WH-V113 belongs to a novel sequence type ST2058 with an allelic profile of *adk*(24), *gyrB*(5), *mdh*(262), *metE*(546), *pntA*(14), *purM*(18), and *pyrC*(45). The *tet*(X4)^G568A^ along with a diverse array of other ARGs—including *tet*(A), *sul2, floR*, and *bla*_OXA-1_ was localized on Chromosome I. This chromosomal integration of the *tet*(X4)^G568A^ variant contrasts sharply with the predominant plasmid-borne localization typically observed in Enterobacterales(3, 4, 16, 17). From an evolutionary perspective, chromosomal integration may stabilize the *tet*(X4) gene through vertical inheritance under fluctuating selective pressures, whereas plasmid carriage typically promotes rapid horizontal dissemination among diverse hosts(18).

To investigate the evolutionary origin of the resistance determinant, we initially assembled 17,518 *V. cholerae* genomes. After using Snippy to filter out sequences with inadequate reference single nucleotide polymorphism (SNP) alignments, yielding a final set of 2,658 genomes for phylogenetic analysis. Population structure analysis via fastBAPS(19) generated 11 distinct clusters **(Fig. 1D)**. Notably, the *tet*(X4)-positive isolate WH-V113 identified in this study localized to cluster BAPS7. To further elucidate its phylogenetic relationships within this specific genetic background, a core-genome SNP tree was constructed using all BAPS7 isolates **(Fig. 1E)**. WH-V113 exhibited the closest phylogenetic affinity to a porcine isolate recovered from China. The remaining isolates in the lineage originated from diverse sources—environmental, clinical, seafood-associated, and unidentified—and were distributed across multiple countries, predominantly in Asia. The absence of phylogenetic clustering associated with host or geography suggests that acquisition of *tet*(X4) represents a recent horizontal gene transfer event rather than inheritance within a stable *Vibrio* lineage.

Detailed mapping of the genetic environment provided direct evidence of the hypothesis. The *tet*(X4)^G568A^ variant was embedded within a highly conserved structural module (IS*CR2* – *tet*(X4) – *abh* – *hp* – *hp-*IS*CR2*) flanked by two repeated IS*CR2*(20, 21) **(Fig. 2A)**. This module exhibits 100% nucleotide identity with the corresponding region of porcine *E. coli* IncX1_1 plasmid pAMR1125_tetX4 (AP050507.1) **(Fig. 2B)**. However, *tet*(X4)-bearing module in WH-V113 was integrated into SXT/R391-like integrative and conjugative element (ICE) at the chromosomal *prfC* locus rather than residing on a plasmid. The nearly identical genetic environments observed in both the *E. coli* plasmid and the *V. cholerae* chromosome strongly suggest that this resistance determinant was recently acquired via plasmid conjugation from an Enterobacteriaceae donor, rather than evolving independently after its introduction into *Vibrio* spp. Furthermore, the configuration of IS*CR2* elements in WH-V113 is capable of forming a classical rolling-circle transposable unit **(Fig. 2C)**. Inverse PCR amplified a 4607bp circular intermediate and sequencing confirmed the excised IS*CR2*–*tet*(X4)–*abh*–*hp-hp* structure **(Fig. 2D)**. These results provide direct experimental evidence that IS*CR2* mediates excision of the *tet*(X4) module through rolling-circle transposition, a mechanism previously proposed but rarely demonstrated experimentally for naturally occurring *tet*(X4) elements(5, 22). Following its transfer, the circular intermediate integrated into the ICE **(Fig. 2E)**. This reveals that *tet*(X4) can be disseminated through a tripartite, complementary mechanism involving IS*CR2*-mediated transposition, plasmid conjugation, and ICE-mediated chromosomal transfer **(Fig. 2E)**. Such profound genetic plasticity greatly enhances the capacity of *tet*(X4) to spread across species barriers and distinct ecological niches, particularly within the food chain(23, 24) **(Fig. 2E)**. The structural conservation further suggests that IS*CR2*-associated modules retain the potential for future remobilization under antibiotic selection, linking long-term chromosomal persistence with renewed horizontal spread.

**Figure 2.**
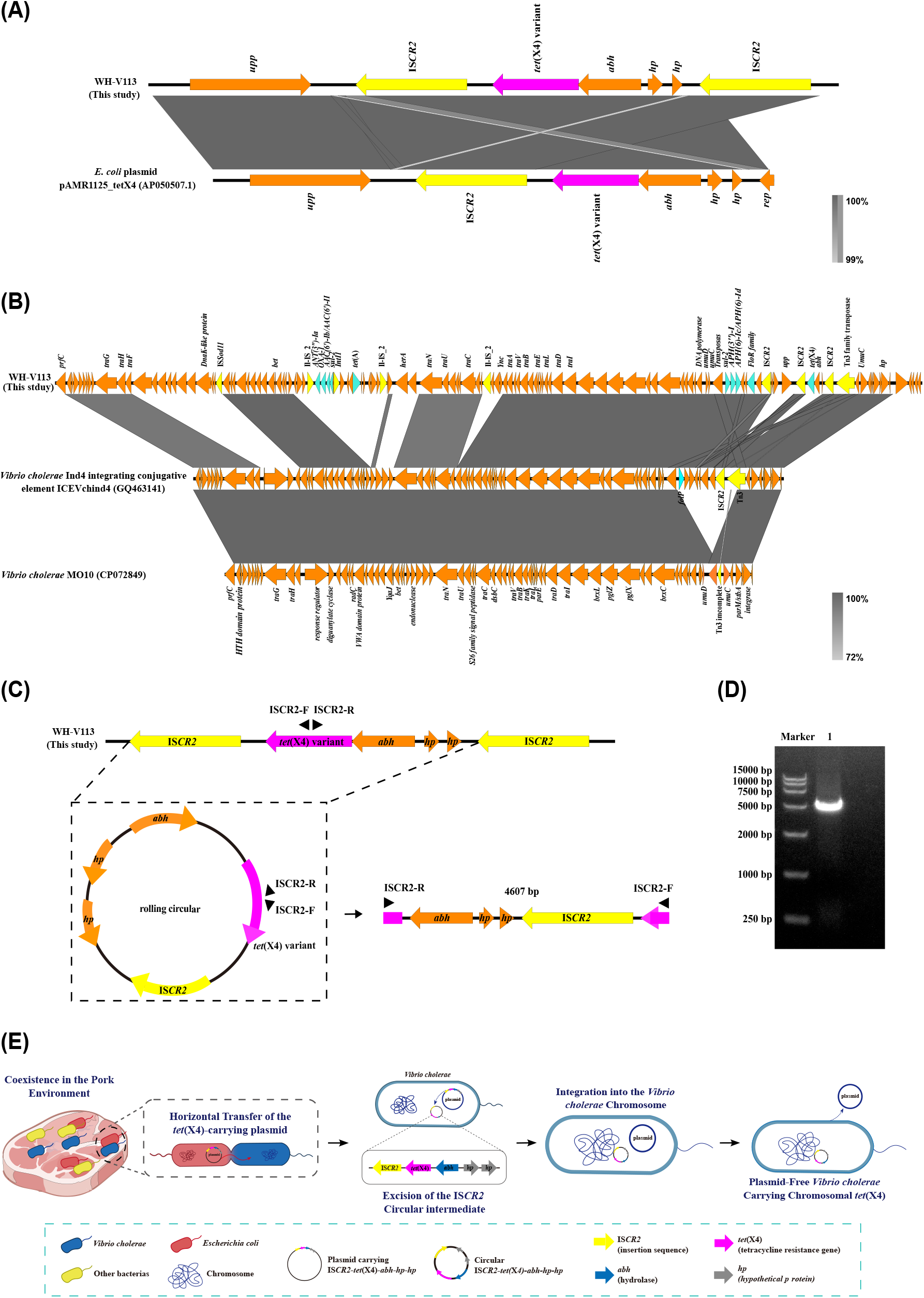
Genetic environment and structural characterization of the *tet*(X4) module. **(A)** Alignment of the conserved *upp*–IS*CR2*–*tet*(X4)–*abh-hp-hp* region among representative sequences with high nucleotide homology. **(B)** Schematic diagram of the inverse PCR strategy. Divergent primers (ISCR2-F/R) were designed on the *tet*(X4) gene to amplify the rolling-circle structure intermediate. **(C)** Agarose gel electrophoresis of the inverse PCR products, showing an expected band of approximately 5kb. **(D)** Alignment of the WH-V113 genomic region with the backbone sequence of the SXT/R391 family integrative and ICE to elucidate the structural arrangement of the *tet*(X4)-carrying ICE. **(E)** Schematic illustrating the mechanism by which the rolling-circle structure IS*CR2*-*tet*(X4)-*abh*-*hp*-*hp* transfers from *E. coli* plasmids to *V. cholerae*.

## Conclusion

Together, our findings demonstrate that a tigecycline resistance determinant predominantly circulating in *Enterobacterales* has entered the environmental *Vibrio* gene pool. The identification of a naturally occurring *tet*(X4)^G568A^ variant further expands the known diversity and host range of *tet*(X4), while highlighting aquatic *Vibrio* species as potential environmental reservoirs and recipients of clinically important antimicrobial resistance genes. Continuous genomic surveillance of environmental *Vibrio* populations is therefore warranted to monitor the emergence and dissemination of mobile tigecycline resistance determinants.

## Acknowledgements

The study was supported by grants from the National Natural Science Foundation of China (no. 32202987, 32300156, and 82572580), the Key Program of Anhui Educational Committee (no. 2022AH050179), Outstanding Innovative Research Team for Molecular Enzymology and Detection in Anhui Provincial Universities (2022AH010012), and Innovation and Entrepreneurship Training Program for college students of Anhui Normal University.

